# Acquiring thin histological sections of açaí (*Euterpe oleracea* Mart.) seeds for molecular mapping by mass spectrometry imaging

**DOI:** 10.1101/2022.08.20.504526

**Authors:** Felipe Lopes Brum, Gabriel Rocha Martins, Ronaldo Mohana-Borges, Ayla Sant’Ana da Silva

## Abstract

**RATIONALE:** Matrix-assisted laser desorption/ionization imaging mass spectrometry (MALDI-IMS) of tissues became popular in the last decade. Consequently, adapting sample preparation methods for different materials turned to be a pivotal step for a successful analysis, due to the requirement of samples slices of 12-20 μm thickness. However, preparing thin sections compatible with MALDI-IMS for unusual samples is challenging, as existing histological protocols may not be suitable, thus requiring new methods. Açaí (*Euterpe oleracea* Mart.) seed is an example of a challenging material due to its toughness and resistance to crack. Therefore, our goal was to develop a methodology to obtain thin (<20 μm) and entire longitudinal sections of açaí seeds for MALDI-IMS analysis.

**METHODS:** Different strategies were evaluated for obtaining thin cuts of seeds, being the combination of the following steps the most suitable option: (i) softening of seeds by water immersion for 24 h; (ii) longitudinal cut of seeds to obtain half-seeds using a razor blade and a hammer; (iii) half-seeds imbibition in gelatin; (iv) sectioning using a cryostat at −20 °C to obtain samples with <20 μm thickness; and (v) collection of samples in an indium tin oxide coated glass slide covered by a double-face copper tape to avoid sample wrapping and ensure adhesion after unfreezing. (iv) storage of samples in a −80 °C freezer, if necessary.

**RESULTS:** As a result, this adapted sample preparation method enabled the analysis of açaí seeds by MALDI-IMS providing spatial distribution of carbohydrates in the endosperm.

**CONCLUSIONS:** The adaptations developed for sample preparation will help investigate the metabolic and physiological properties of açaí seeds in future studies.

## 1. INTRODUCTION

The açaí palm (*Euterpe oleracea* Mart.) is native to the Amazon region, whose fruit’s pulp consumption and commercialization are expanding in domestic and international markets driven by the claims of its health benefits^1,2^. In 2020, Brazil produced almost 1.7 million tons of açaí^3^. However, açaí edible part represents only 15% of the mature fruits’ wet weight^4^, whereas the açaí seed accounts for the other 85%. Thus, almost 1.4 million tons of seeds disposed of as agro-industrial waste yearly are discarded as consequence of pulp production, and only a tiny amount of this residue is currently exploited as a co-product in other industries^3,5,6^. As Brazil is the primary açaí producer globally, the seeds’ disposal represents a significant environmental, sanitary, and economic issue^6–8^.

Nevertheless, previous reports show that the seeds are a source of bioactive compounds with the potential for high-value bioproducts production from this residue^5,9–12^. Therefore, studies focusing on describing the composition and properties of açaí seeds can provide information to help implement strategies for its uses and industrial applications.

Morphologically, the mature açaí seed is enveloped by a fibrous mesocarp and formed by an endocarp that contains a tiny embryo, an abundant endosperm with resistant properties, and a ruminated tegument^4,13–15^. Identifying the location and content of specific compounds across a tissue may help correlate to their function^16^. However, there is a very small amount of information about the localization of primary compounds in açaí seeds. A previous study has quantitatively analyzed the proteome dynamics and morphological aspects of the açaí pericarp and seed during the developmental stages^17,18^. Also, a recent histochemical study provided the presence of phenolic compounds and carbohydrates in the açaí seed coat and endosperm, respectivelly^19^. However, knowing its compounds’ distribution and specific composition can help develop optimized processing strategies to produce new products aiming to upgrade the açaí productive chain.

In this context, matrix-assisted laser desorption/ionization imaging mass spectrometry (MALDI-IMS), a technique commonly used in other fields, like medical diagnosis or pharmaceutical research^20,21^, can provide this information if an adequate sample preparation is performed. MALDI-IMS combines the high-resolution identification of thousands of compounds with the information on their spatial distribution directly on the surfaces of the analyzed tissue^22–25^. In contrast, microscopy techniques can only identify a few molecules of interest by staining or immunostaining^26^.

MALDI-IMS efficiency depends on the ionization properties of an analyte and appropriate matrix rather than on staining or antibody affinity of few target compounds. In MALDI-IMS analysis, the sample slice must be placed on a glass slide covered by indium tin oxide (ITO) or other conductive surface that favors the ionization step. The slice’s thickness and flat surface are fundamental for the start point desorption^27^. However, it is still challenging to prepare thin sections of unusual samples to be compatible with MALDI-IMS using existing histological protocols, thus requiring adaptations. Therefore, improving sample preparation protocols has become essential to obtain better results with this technique. In the case of açaí seeds, the challenge of its preparation for MALDI-IMS analysis relies on its toughness and resistance to crack, as açaí seed can withstand mechanical damage even when submitted to forces as high as 950 N due to the resistance properties of linear mannan, a reserve polysaccharide found in the seed’s endosperm^11^.

For this reason, physical fixation, such as cryofixation, which is broadly used as a preparation method to obtain thin sections of samples^25^, is not appropriate for this particular seed because the standard protocol mainly contemplates soft tissues. Therefore, for a successful MALDI-IMS analysis, sample preparation becomes a pivotal part of the method.

Recently, our group published experimental data on polyphenols analysis of the açaí seed by MALDI-IMS^10^. However, challenges regarding the sample preparation method were not the focus of that work and, therefore, were not described and discussed in detail. Thus, this work presents in detail the adapted protocol for preparing açaí seed samples to be analyzed appropriately by MALDI-IMS and shows its adequacy to map carbohydrates across the açaí seed endosperm, aiming to help other research groups with similar difficulties in preparing resistant materials.

## 2. MATERIAL AND METHODS

### 2.1 Materials

A single-edge carbon steel razor blade Cat.#71960 (Hatfield, PA, USA) was used for longitudinal cut to obtain half-seeds. Gelatin from bovine skin (G9382), carboxymethylcellulose (C4146), 2,5-dihydroxybenzoic acid (DHB, 98%), acetone and acetonitrile were purchased from Sigma Aldrich (St. Louis, MO, USA). A Glass indium tin oxide (ITO) covered slides, and optimal cutting temperature compound (OCT) were purchased from Fisher Scientific (Kalamazoo, MI, USA). Other materials were purchased locally in Rio de Janeiro (Brazil), such as silicon ice mold, aluminum foil, paraffin wax, standard and copper double-faced adhesive tapes, and superglue from Loctite (1621047 Henkel, Brazil).

### 2.2 Botanical sample

Açaí Amazonas Ltd. (Óbidos, Pará, Brazil) kindly donated the açaí seeds obtained after the industrial depulping process. The seeds were kept in sealed boxes at room temperature for further analysis. Optical images of the seeds’ sections were taken in a stereomicroscope (SteREO Discovery.V8, Carl Zeiss, 172 Germany) with the support of a cell phone camera coupled with a 3D-printed cell phone adapter for a microscope (EMCUBOS, Rio de Janeiro, RJ, Brazil).

### 2.3 Developing a sectioning protocol

The açaí mesocarpic fibers layer was manually removed to better adhere the seed to the holder of the microtome or cryostat and to improve the interaction with all inclusion media tasted.

Different histological methods were adapted for MALDI-IMS optimal spectra acquisition to ensure sample thickness (10-20 μm). Manual sectioning using a razor blade or sectioning assisted by equipment were evaluated (microtome and cryostat) to obtain thin, uniform, and flat sections of the açaí seeds. The equipment-assisted sectioning process used a rotary microtome (820 Spencer Microtome, American Optical Corporation, USA) and a cryostat (Leica CM1860-UV, Leica Biosystems, Nussloch, Germany). Prior to sectioning trials, different strategies to prepare the sample were tested including: samples’ water immersion time (up to 7 days), fast freezing of *in natura* samples (by immersion in liquid nitrogen, inside a freezer at −80 °C or immersion in a mixture of acetone and dry ice) and evaluation of various embedding media to facilitate cutting (paraffin, carboxymethyl cellulose, and gelatin). After the suitable cut was obtained, standard and copper double-faced adhesive tapes were tested to guarantee adhesion to glass slides.

### 2.4 Matrix-assisted laser desorption/ionization imaging mass spectrometry (MALDI-IMS) experimental procedures

Before MALDI-IMS analysis, a dihydroxybenzoic acid (DHB) matrix was added to the samples using a standard sublimation method ^28,29^. Afterward, the samples were scanned, and the spectra were acquired in positive mode with a 75 μm lateral resolution, 200,000 mass resolution from 150 to 3000 Da in a Solarix XR FT-ICR 7T (Bruker Daltonics, Germany) spectrometer with a nitrogen laser (337 nm), and the data was processed using the SCiLS Lab MVS Version 2019c Premium 3D software.

### 2.5 Elementary composition analysis by EDS

The açaí seed samples were cut using an IsoMet 1000 Precision Cutter (Buehler, Lake Bluff, IL, USA) equipped with an IsoMet Diamond Wafering Blade, 15HC (Buehler, Lake Bluff, IL, USA), to obtain a 2.0 mm slice section that was fixed in the stub with carbon tape. Samples were analyzed for their elementary composition and surface morphology, performed in a scanning electron microscope Inspect S50 (FEI Company, Hillsboro, OR, USA), associated with an X-ray energy dispersive spectrometer (EDS) SDD type, Apollo 10 model (EDAX, Pleasanton, CA, USA). The images were acquired at 20 kV with spot-size 7.0 over a working distance (WD) equal to 10 mm, with a backscattered electrode detector (BSED) in a low vacuum (50 Pa). EDS spectra were acquired under the same voltage and spot conditions previously mentioned, with signal strength greater than 3000 counts per second (CPS) and dead time (DT) between 20 and 30 percent. In these same conditions, elementary maps were acquired, at a resolution of 128×128 pixels, with an accumulation of 32 frames.

## 3. Results and Discussion

### 3.1 Sample preparation method

MALDI-IMS analysis requires very thin slices of tissue (12-20 μm)^30^ with a flat surface, which are fundamental for an efficient ionization and desorption^27^. However, açaí seeds (**Figure 1a**) present a particular toughness, probably associated with its high content of linear mannan in the endosperm (**Figure 1b - x**), a crystalline polysaccharide^31^, and the high content of procyanidins in a thick external tegument layer (**Figure 1b**), that is reported to contribute to seed coat toughness ^10,32,33^. Thus, routine sample preparation methods for MALDI-IMS analysis were incompatible with the seed’s attributes, requiring the development of an adapted protocol, as its resistance to cut hampered the acquisition of thin and entire sections of the sample using traditional preparation methods such as direct sectioning with a sharp blade, followed by samples’ imprinting, or fixation with organic solvents ^25,27^.

**Figure 1.**
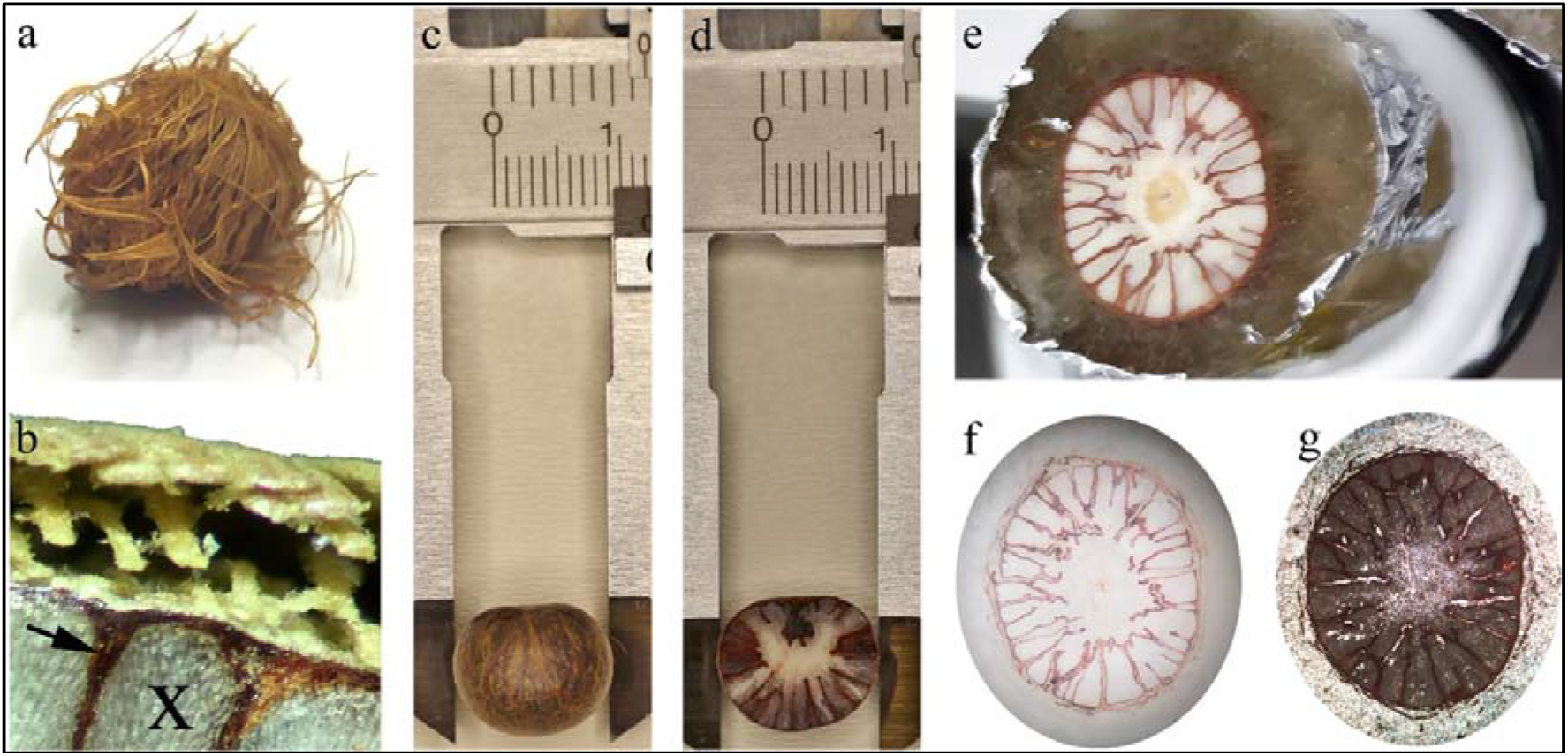
Sample preparation of açaí seed for MALDI-IMS analysis. a) Açaí seed with mesocarpic fibers. b) Inset of the seed sectioned in half, with identifiable mesocarpic fibers (upper part), endosperm (x) and tegument projections (arrow). c) The wet seed without external fibers d) A half-cut wet seed without external fibers. e) A half-cut wet seed without external fibers embedded in gelatin (transparent) already sectioned inside the cryostat; the OCT (white) just below the gelatin; and the side of the holder (black) just below the OCT. f) A seed slice adhered in double-sided tape. g) A seed slice adhered in double-sided copper tape. c-d were measured by a precise measuring instrument (Zaas). f-g images were obtained with a stereomicroscope (SteREO Discovery.V8, Carl Zeiss, 172 Germany).

Before evaluating preparation protocols, the mesocarpic fibers that cover the açaí seed (**Figure 1a and b – upper part**) were removed (**Figure 1c**). The first attempt to section açaí seed *in natura* with a sharp razor blade resulted in the complete sample fracture even when individualized samples were previously frozen by immersion in in liquid nitrogen, or in a mixture of acetone and dry ice or directly frozen at −80 °C. Then, a new effort was made using a microtome with the sample adhered to the holder with superglue. However, due to the seed’s resistance to crack, the superglue was insufficient to hold the globose seed when the steel blade forced its surface, and the sample dropped from the holder. Next, a strategy to use embedding media to facilitate sectioning with a microtome was performed, as previous studies with tooth^34^, craniofacial pig bone^35^, and botanical samples^36–38^ successfully applied this method. Paraffin was first evaluated, as it is reported to facilitate sectioning using a microtome at room temperature. However, the paraffin embedding could not allow the seed sectioning, resulting in fragmentation of the sample and block breakage.

Therefore, we applied other embedding media (carboxymethyl cellulose (CMC) and gelatin) and a cryostat (subzero) to attempt the sample sectioning. In a cryostat acclimatized to −20 °C, the sample was placed in a plastic collector and submerged in a mixture of acetone with carbon dioxide in a solid-state (dry ice), aiming the ideal freezing for the subsequent preservation at −80 °C. We used OCT to adhere the sample to the cryostat holder. However, during sectioning trials, only dispersed and fragmented seed samples were obtained.

Then, before applying the embedding media, we evaluated immersing the sample in deionized water for 1, 3, 24, 72, and 168 h, aiming to soften the seeds. Then, a longitudinal cut was made to obtain half-seeds with a razor blade and hammer, and two different media were used to embed the samples, CMC (2-4% in water) or gelatin (100 mg/mL in water), aiming to obtain a whole seed slice in the cryostat (**Figure 1e**), repeating the freezing procedure described above. The CMC medium was inefficient in facilitating sample sectioning, resulting in sample crack. However, gelatin embedding resulted in sectioning of almost integral seed’s slices with proper thickness (~20 μm) when a minimum of 24 h of water-immersed seeds were evaluated **(Figure 1c-d)** (1 and 3 hours did not result in efficient sectioning inside the cryostat).

However, immediately after sectioning, the sample curled similarly to pencil shavings. This property hampered adhesion to the glass slides covered with indium tin oxide (ITO) in a flat format. Then, a double-sided adhesive tape, **Figure 1f**, was used to help adhesion^39^, but the slices lost adhesion after unfreezing from the storage at −80 °C. As a solution, a double-sided copper tape provided better adhesion to the glass slide besides increasing the conductivity and optimizing ionization (**Figure 1g**).

**Figure 2** summarizes the steps that resulted in preparing thin (<20 μm) sections of açaí seeds. We suggest the following protocol for sample preparation:

1. remove the fibrous mesocarp;
2. soak the sample in water for at least 24 h;
3. cut the seed with a razor blade and hammer longitudinally;
4. put the wet sample in freezable plastic, silicon, or aluminum foil mold and add fresh-made warm gelatin medium (100 mg/mL in water);
5. submerge this mold in a mixture of acetone with carbon dioxide in a solid-state (dry ice), taking care not to allow the mixture to come into direct contact with the gelatin or the sample;
6. put the sample inside a cryostat acclimatized to −20°C and wait for 2 h;
7. collect the sample slices at 20 μm of thickness with ITO coated glass slide covered by a double-face copper tape, then preserve at −80°C if necessary;
8. sublimate the MALDI-IMS chosen matrix;
9. generate an image of the entire glass slide by scanning;
10. put the glass slide inside the proper image spectrometer;
11. use SCiLS Lab or other appropriate software to convert the data acquired as spectral graphs into image.

**Figure 2.**
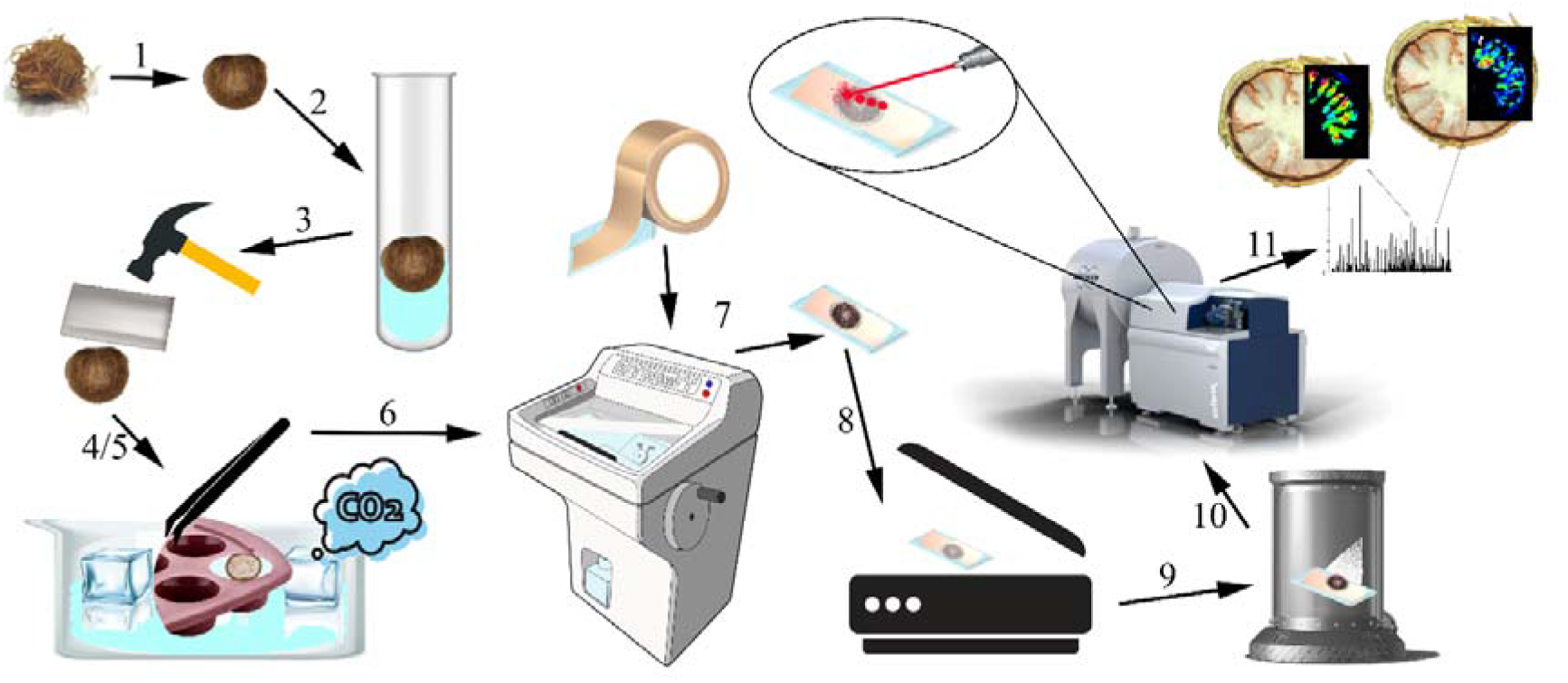
Schematic representation of the seed processing method steps for IMS analysis. The numbers are related to the description of the steps in the text (Images: Adobe Stock*/Illustration: Felipe L. Brum). *All images used have free licenses or were provided by the authors.

### 3.2 MALDI-IMS analysis using the sample prepared by the adapted protocol

To validate the sample preparation method, seeds slices were analyzed aiming to map the distribution of carbohydrates in the seeds. The compound identification using MALDI-IMS was assisted by previous studies of the *E. oleracea* Mart. mature seed. Açaí seed is formed by the seed coat, called testa or tegument, an embryo, and the endosperm^13,19^. The endosperm is majorly composed of mannan (**Figure 1b, x**), a polymer of the hexose mannose, which provides embryo protection, and serves as a reserve for seedling nutrition and development^11,14^. The MALDI-IMS analysis exhibited peaks representing [M+K]^+^ adducts of hexose oligomers (Δ= 162 Da), providing molecular information at the tissue level (**Figure 3**). Hexose dimers (*m/z* 381), trimers (*m/z* 543), and tetramers (*m/z* 705) appeared almost exclusively in the core of the endosperm. From the hexamer (*m/z* 1029) up to a fourteen-unit oligomer (*m/z* 2325), these carbohydrates appear in the outer tissues, in the endosperm near the testa ruminations, although. they seem to be absent in the tegument projections. This could indicate that, besides accumulating mannan with high degree of polymerization (DP), low DP oligomers are readily available in the mature and dormant seed to trigger the germination process.

**Figure 3.**
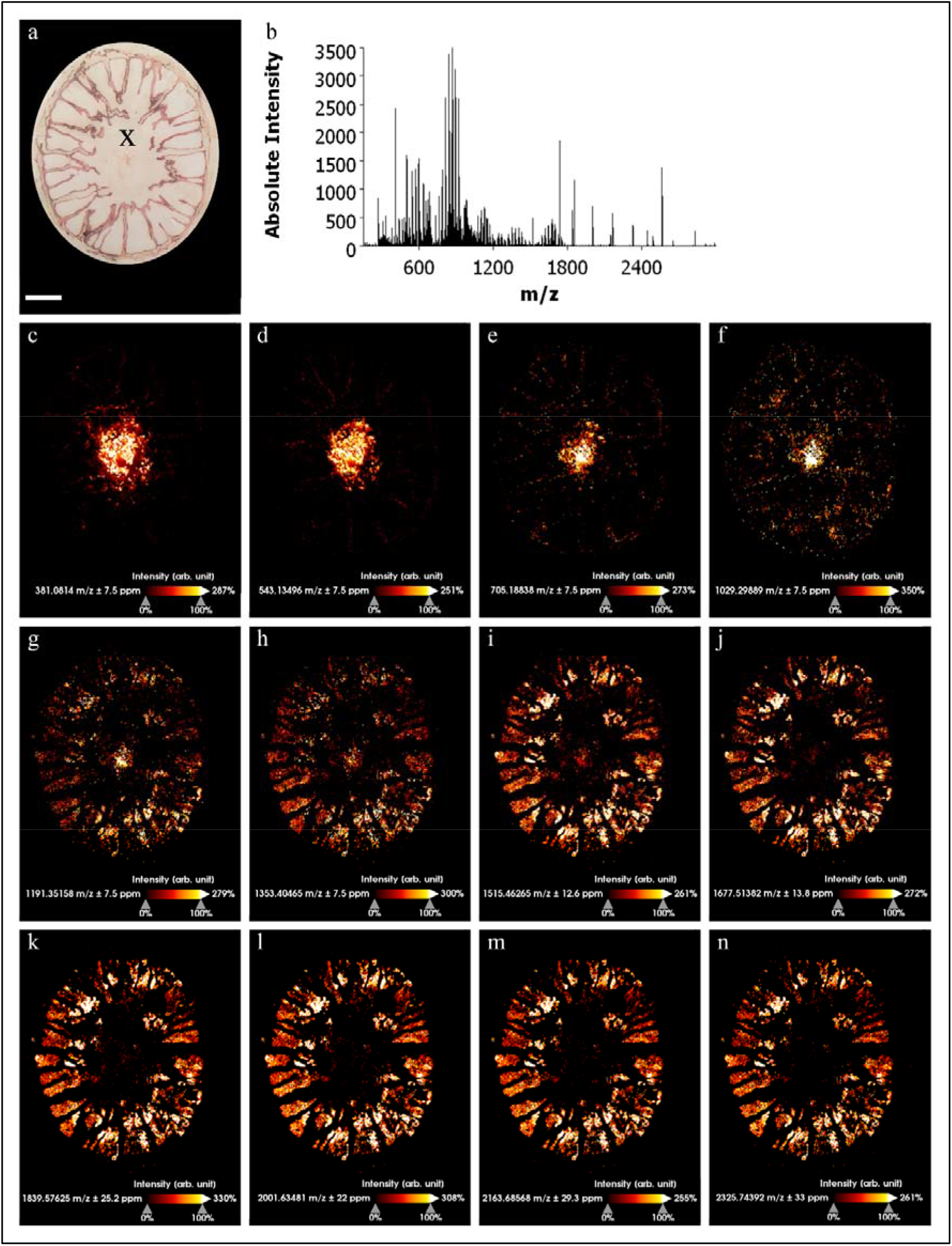
Distribution of hexose oligomers in açaí seed by MALDI-IMS in positive mode. a) Histology of the seed slice analyzed pointing the center of the endosperm (x). Scale bar = 2mm.. b) Spectra showing all molecular peaks acquired. c-n) molecular image of the hexose oligomers distribution.

As mentioned, MALDI-IMS data exhibited *m/z* peaks as potassium adducts without previously adding salt to the matrix during the deposition step. Thus, to understand if this was an artifact introduced by the adapted sample preparation method or a characteristic of the sample, energy dispersive X-ray spectrometry (EDS) was used for qualitative identification and semi-quantitative elemental information of the açaí seed^40^. The EDS analysis **(Figure 4)** showed that the elements detected were ubiquitous and uniformly distributed in both analyzed tissues (endosperm and tegument). The organic fraction, represented by the non-mineral elements C and O was significantly higher than the minerals found, mostly associated with carbohydrates. Potassium (K) was the most accumulated mineral element corresponding to 6.25% (w/w) of the seed, thus confirming that potassium adducts were formed during the MALDI-IMS analysis due to samples’ characteristics and not as an introduced artifact. Therefore, the adapted sample preparation method showed to be suitable for mapping the spatial distribution of molecules in the açaí seed.

**Figure 4.**
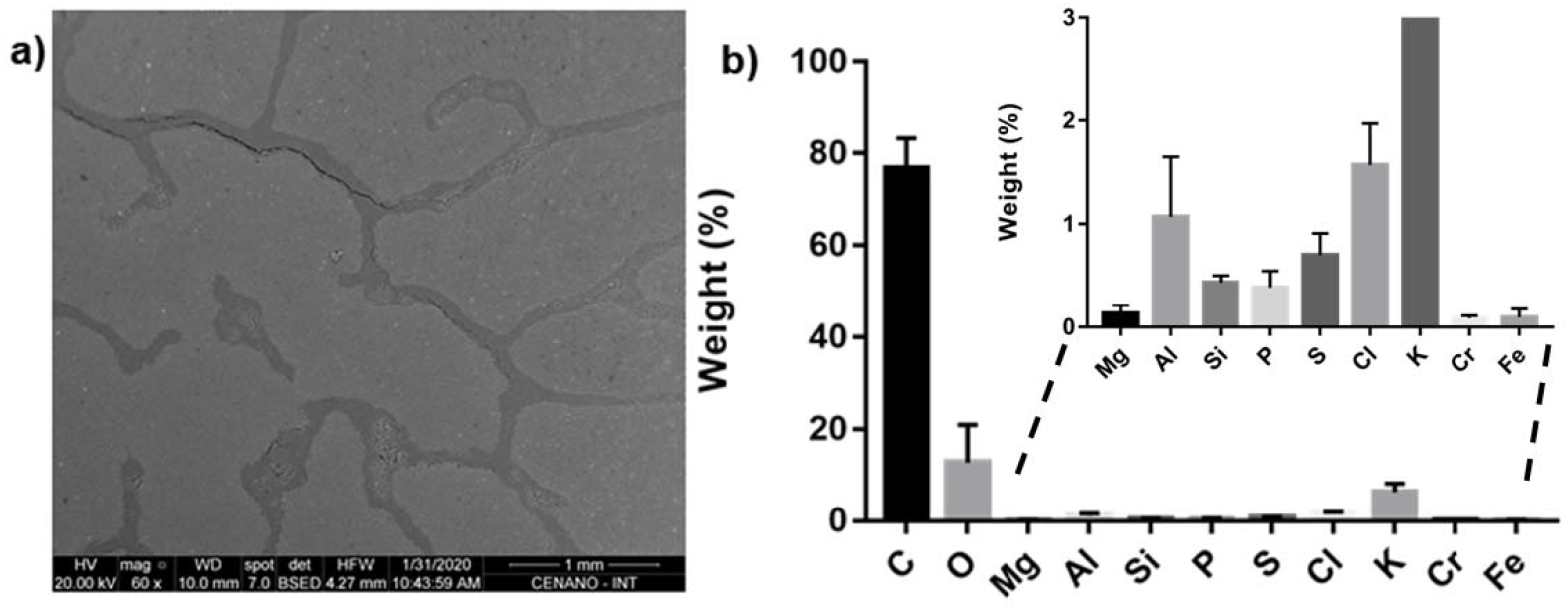
Elementary composition analysis of açaí seed by energy dispersive X-ray spectrometry. a) Açaí seed sample section. b) The main elements quantified. The elemental abundance corresponds to about 76.6% of carbon (C), 12.7% of oxygen (O), 6.25% of potassium (K) and less than 5% of the others (Mg, Al, Si, P, S, Cl, Cr and Fe).

## 4. CONCLUSIONS

This study provided an approach for preparing açaí seeds for MALDI-IMS for mapping metabolites. The sample preparation protocol devised enabled the analysis of with excellent resolution, helping the identification of spatial distribution of metabolite composition. The adapted protocol was designed based on different preparation methods, tailored for a MALDI-IMS analysis. In the future, this technique may be used to investigate physiological and molecular mechanisms during initial stages of açaí seed germination or final stages of seed development, providing more comprehension of the metabolomic changes during these phenomena.

## ACKNOWLEDGMENTS

This work was financed by Serrapilheira Institute (Serra-1708-15009) and the Coordination for the Improvement of Higher Education Personnel (CAPES-AUXPE 0415/2016) and Carlos Chagas Filho Foundation for Supporting Research in the State of Rio de Janeiro (FAPERJ-JCNE-SEI-260003/004754/2021). Serrapilheira Institute and the National Council for Scientific and Technological Development (CNPq) granted scholarships for Dr. Felipe Lopes Brum and Dr. Gabriel R. Martins (Institutional Capacity Building Program/INT/MCTI). Centro de Espectrometria de Massas de Biomoleculas (CEMBIO-UFRJ) is acknowledged for the services provided with MALDI-IMS analyses, and the Centro de Caracterização em Nanotecnologia para Materiais e Catálise (CENANO-INT) funded by MCTI/SISNANO/INT-CENANO-CNPQ grant N° 442604/2019 is thanked for the elementary composition analysis.

## CONFLICT OF INTEREST STATEMENT

The authors declare no conflict of interest.

## CREDIT AUTHORSHIP CONTRIBUTION STATEMENT

**FLB**: Conceptualization, Data curation, Formal analysis, Investigation, Methodology, Roles/Writing - original draft. **GRM**: Conceptualization, Data curation, Investigation, Roles/Writing - original draft. **RMB**: Methodology, Software, Supervision, Writing - review & editing. **ASS**: Funding acquisition, Investigation, Project administration, Resources, Supervision, Writing - review & editing.

## DATA AVAILABILITY STATEMENT

The data that support the findings of this study are available on request from the corresponding author.

## Notes

Funding Information: Serrapilheira Institute, grant number Serra-1708-15009; Coordination for the Improvement of Higher Education Personnel (CAPES), grant number AUXPE 0415/2016; Carlos Chagas Filho Foundation for Supporting Research in the State of Rio de Janeiro (FAPERJ-JCNE-SEI-260003/004754/2021).

### Competing Interest Statement

The authors have declared no competing interest.

